# Genetic differences in the behavioral organization of binge eating, conditioned food reward, and compulsive-like eating in C57BL/6J and DBA/2J strains

**DOI:** 10.1101/190827

**Authors:** Richard K. Babbs, Julia C. Kelliher, Julia L. Scotellaro, Kimberly P. Luttik, Megan K. Mulligan, Camron D. Bryant

## Abstract

Binge eating (**BE**) is a heritable symptom of eating disorders associated with anxiety, depression, malnutrition, and obesity. Genetic analysis of BE could facilitate therapeutic discovery. We used an intermittent, limited access BE paradigm involving sweetened palatable food (**PF**) to examine genetic differences in BE, conditioned food reward, and compulsive-like eating between C57BL/6J (**B6J**) and DBA/2J (**D2J**) inbred mouse strains. D2J mice showed a robust escalation in intake and conditioned place preference for the PF-paired side. D2J mice also showed a unique style of compulsive-like eating in the light/dark conflict test where they rapidly hoarded and consumed PF in the preferred unlit environment. BE and compulsive-like eating exhibited narrow-sense heritability estimates between 56 and 73 percent. To gain insight into the genetic basis, we phenotyped and genotyped a small cohort of 133 B6J × D2J-F_2_ mice at the peak location of three quantitative trait loci (**QTL**) previously identified in F_2_ mice for sweet taste (chromosome 4: 156 Mb), bitter taste (chromosome 6: 133 Mb) and behavioral sensitivity to drugs of abuse (chromosome 11: 50 Mb). The D2J allele on chromosome 6 was associated with greater PF intake on training days and greater compulsive-like PF intake, but only in males, suggesting that decreased bitter taste may increase BE in males. The D2J allele on chromosome 11 was associated with an increase in final PF intake and slope of escalation across days. Future studies employing larger crosses and genetic reference panels comprising B6J and D2J alleles will identify causal genes and neurobiological mechanisms.

## 1. INTRODUCTION

Eating disorders (**ED**) are the most lethal of all psychiatric disorders, with high mortality rates and lifetime prevalence rates ranging from 1-3% [1]. ED and related behaviors, including binge eating (**BE**) are heritable; however, the genetic basis of ED and BE remain largely unknown [2]. BE can be observed in all three major EDs, including Binge Eating Disorder (**BED**), Bulimia Nervosa (**BN**), and Anorexia Nervosa (**AN**) [3, 4]. BE is defined by the compulsive/unrestrained consumption of an extraordinarily large amount of food (typically palatable food with a high caloric content) within a brief time period and is accompanied by a sense of loss of control as well as feelings of uncomfortable fullness, distress, disgust, anxiety, depression, guilt, and remorse [5]. BE is also associated with hoarding behavior [6], including hoarding of food so that it can be consumed alone [7]. BE is associated with multiple health issues, including mood dysfunction, aberrant eating patterns and cycles of BE and food restriction, malnutrition, and obesity-related health issues [8]. BE can be extinguished in humans and in preclinical models [9, 10]; however, there is a high rate of rapid and long-term relapse [11, 12].

Potential pharmacotherapeutic treatment options for BE have had limited success. The only FDA-approved drug for the treatment of moderate-to-severe BED is Lisdexamfetamine dimesylate, an amphetamine prodrug with limited, long-term efficacy and a high risk of abuse potential and adverse cardiovascular events [13, 14]. Accordingly, treating BE-presenting patients with amphetamines could be problematic, given the high degree of comorbidity and shared neurobiological mechanisms between BE and substance use disorders [15, 16]. Clearly, more efficacious treatments with fewer side effects are needed.

Very few genome-wide forward genetic (linkage or GWAS) studies have been conducted for BE or BED in humans [17] or in preclinical models [18]. In rodents, BE in non-stressed, non-food-deprived animals is most readily achieved under a specific set of conditions that includes 1) presentation of palatable food (e.g., sweetened or high fat), 2) limited access (e.g., 0-2 h), and 3) intermittent access [19, 20]. We recently utilized these conditions to conduct a genome-wide quantitative trait locus (**QTL**) study of BE between closely related C57BL/6 substrains and mapped and validated cytoplasmic FMR1-interacting protein 2 (*Cyfip2*) as a major genetic factor underlying BE [18]. Owing to the drastically reduced genetic complexity of this cross, BE exhibited a simple Mendelian inheritance, yielding a single locus explaining the parental strain (genetic) variance in BE [18]. This highlights both a main advantage and disadvantage of utilizing reduced complexity crosses for forward genetic studies of complex traits. One the one hand, reduced genetic complexity can quickly lead to identification of the underlying genetic factors responsible for behavioral trait variation [18, 21–23]. On the other hand, complex traits in a reduced complexity cross are expected to segregate phenotypic variance in a near-Mendelian fashion and thus, only one major locus/gene is likely to account for most of the genetic variation in a complex trait [18, 22, 23]. Ultimately, populations of increasing complexity must be employed to understand the genetic basis of complex traits such as BE.

C57BL/6J (B6J) and DBA/2J (D2J) are two of the most widely used inbred strains in forward genetic studies of complex traits, largely because they are the original parental strains of one of the largest and most widely used set of murine recombinant inbred (**RI**) strains for dissecting the genetic basis of neurobehavioral traits ‑ the BXD RI panel [24]. A heritable strain difference in BE and associated behaviors between B6J and D2J provides the foundation for future use of the BXD RI panel in a systems genetics approach to advance our knowledge regarding the preclinical genetic and neurobiological basis of BE and ultimately translate these findings to humans.

To facilitate the long-term goal of conducting genetic mapping studies in additional crosses and panels, in the present study, we assessed strain differences in BE of sweetened palatable food (**PF**) and concomitant behaviors between the BE-resistant B6J strain [18, 25] and the D2J strain which has previously been shown to exhibit either less or more consumption of sweetened solutions relative to the B6J strain, depending on the duration of access and the concentration of solution [26–31]. We next estimated narrow-sense heritability of 16 BE-related traits. To provide further evidence for a genetic basis for BE, we phenotyped a small cohort of 133 B6JxD2J-F_2_ mice (**F_2_**) and genotyped these mice at markers targeting these three QTLs, including a first QTL associated with sweet taste (chromosome 4: 156 Mb) [26, 31, 32]), a second QTL associated with bitter taste (chromosome 6: 133 Mb) [26, 33], and a third QTL associated with behavioral sensitivity to methamphetamine and cocaine (chromosome 11: 50 Mb) [34–36].

## 2. METHODS AND MATERIALS

### 2.1. Mice

All experiments were conducted in accordance with the NIH Guidelines for the Use of Laboratory Animals and were approved by the Institutional Animal Care and Use Committee at Boston University. Mice were maintained on a 12 h /12 h light/dark cycle (lights on at 0630 h) and housed two to four animals per cage in same-sex cages. Laboratory chow (Teklad 18% Protein Diet, Envigo, Indianapolis, IN, USA) and tap water were available *ad libitum* in home cages. Testing was conducted during the light phase. C57BL/6J (**B6J**), DBA/2J (**D2J**) and B6JxD2J-F_1_ mice (7 weeks old) were purchased from the Jackson Laboratory (JAX; Bar Harbor, ME) and were generated at JAX by crossing B6J males with D2J females. Purchased mice were habituated to the vivarium one week before testing in a separate room. B6JxD2J-F_2_ mice were generated in-house by pairing the F_1_ mice. Mice were between 50 and 100 days old on the first day of training.

### 2.2. BE, CPP, and the light/dark conflict test of compulsive-like eating

On Day (D)1, mice were assessed for initial preference for the food-paired side (the right side) by placing the mice into the left side, providing open access (5 cm × 6.5 cm) to both sides for 30 min, and video recording initial time spent on the food-paired side. On D2, D4, D9, D11, D16, and D18, mice were restricted to the right side of the apparatus (pointed floor texture) and given access to 40 sweetened PF pellets (20 mg each, 5TUL diet, Test Diet, MO, USA) or 40 unsweetened Chow pellets (20 mg each, 5BR3 Test Diet, MO, USA) of a similar size and weight for 30 min. In contrast, on D3, D5, D10, D12, D17, and D19, mice were restricted to the left side (smooth-textured floor) with an empty bowl for 30 min. On D22, mice were assessed for final side preference by placing the mice in the left side and providing open access.

On D23, we used the light/dark conflict test to assess compulsive-like eating and concomitant behaviors [18]. Briefly, light/dark boxes had two chambers (20 cm × 40 cm) connected by a small entryway. The dark, unlit, normally preferred chamber had black, opaque sides and a top and bordered the clear, light-exposed, aversive side. A white porcelain bowl containing PF was secured to the middle of the light side floor. Because rodents show a natural aversion for light, compulsive-like eating was operationalized as the intake of PF in the light side [18]. Additional non-ingestive compulsive behaviors were recorded using AnyMaze video tracking software (Stoelting, Wood Dale, IL, USA). Only a subset of data from the Chow-trained D2J group on D23 were available for analysis (n=12/23 total samples).

### 2.3. Behavioral testing and genotyping in B6J × D2J-F_2_ mice

We phenotyped a total of 133 F_2_ mice (68 females, 65 males) in our intermittent, limited access, CPP paradigm. A subset of 93 of these mice (43 females, 50 males), were tested in a battery of premorbid compulsive-and anxiety-like behaviors over one week prior to PF training. F_2_ mice were genotyped for SNPs located at three historic B6J/D2J-derived quantitative trait loci (**QTLs**): a chromosome 4 QTL associated with sweet taste within the *Tas1r3* gene (rs6316711; 156 Mb; [37]), a chromosome 6 QTL associated with bitter taste within the *Tas2r110* gene (rs13479039; 133 Mb; [33]), and a third QTL on chromosome 11 associated with behavioral sensitivity to methamphetamine and cocaine upstream of the *Hnrnph1* gene (rs29383600; 50 Mb) [34–36]. SNP genotyping was performed using fluorescent markers (Life Technologies, Carlsbad, CA USA) and a StepOne machine (Applied Biosystems, Foster City, CA USA).

### 2.4. Premorbid compulsive- and anxiety- like behaviors B6J × D2J-F_2_ mice

Because of the link between anxiety, compulsivity and pathological overeating [38] and because obsessive-compulsive behavior is associated with eating disorders [39, 40], following the identification of parental strain differences in BE, we developed a behavioral battery to assess differences in premorbid anxiety-like and compulsive-like behaviors [41] that we applied toward experimentally naïve F_2_ mice. A subset of the F_2_ mice we generated (92 out of 132 mice total) were tested in the behavioral battery during the first week of testing with one test per day over five days in the following order: 1) open field; 2) elevated plus maze; 3) marble burying; 4) hole board; 5) y-maze. Procedural details are provided in the Supplementary Information. The following week, training commenced for the BE protocol.

### 2.5. Behavioral analysis of BE in B6J, D2J, B6J × D2J-F_1_, and B6J × D2J-F_2_ mice

B6J and D2J mice were tested in a 2 × 2 factorial design to examine PF or Chow intake as a function of Genotype (Strain). Data analysis was conducted in R (https://www.r-project.org/). Food intake (normalized to body weight), CPP (time spent on the food side pre-versus post-training), compulsive-like eating (PF intake on D23 normalized to body weight), and behavior (percent time spent in light, number of entries to light, number of freezing episodes in light, and mean visit time) were analyzed using mixed-model ANOVA (Genotype, Treatment, and Sex as independent variables; Day as a repeated measure). *Post hoc* one-or two-way ANOVA and Welch’s unequal variances *t* tests (p < 0.05, Bonferroni-corrected) were also conducted to determine effects on individual days. Slope analysis of normalized food intake across the food training days was conducted to quantify escalation [18, 25, 41, 42]. Due to a video recording failure caused by insufficient computer storage space, non-ingestive behaviors (time on, number of visits to, and mean visit time on the light side) on D23 were not obtained in 8 of the 16 PF-trained F_1_ mice. Therefore, data from only 8 F_1_ PF-trained mice are reported for the non-ingestive D23 phenotypes.

### 2.6. Heritability estimates of BE and concomitant phenotypes

Narrow-sense heritability (h^2^) was estimated using the between-strain variance (genetic variance; V_G_) and the average within-strain variance (environmental variance; V_E_) of the parental strains and F_1_ mice for each phenotype and then calculating the percentage of the total phenotypic variance (V_G_+V_E_) accounted for by V_G_ (i.e., heritability) according to the following formula: h^2^ = V_G_/(V_G_+V_E_) *100 [43, 44]. This analysis is based on the fact that individuals within a strain are genetically identical and thus, any difference within a strain is assumed to be due to environmental effects and any difference between strains is assumed to be due to genetic differences. This analysis does not account for non-additive variance explained by gene × environment or gene × gene interactions. In cases where we identified statistically significant effects of Genotype, we report percent phenotypic variance accounted for (i.e., h^2^ explained) at the chromosomal locus by dividing the genetic variance between genotypes at the particular locus (marker) by the total phenotypic variance in F_2_ mice. Percent genetic variance explained by a given locus for a trait was calculated by dividing the genetic variance of the locus by the genetic variance of the trait.

### 2.7. Correlation matrices and exploratory factor analysis

To examine the correlation among the major behavioral phenotypes, correlation matrices of variables collected in PF-trained B6J, D2J, and B6J × D2J-F_2_ mice were generated in R using the “rcorr” function from the package “Hmisc.” This function employed Spearman’s rank pairwise correlations across 16 behavioral variables. To reduce the dimensionality of the data and potentially uncover novel relationships, we conducted exploratory factor analysis [45]. This analysis was conducted using the R package “psych” using the “factanal” function. We used the “Varimax” function for orthogonal rotation of the matrix, yielding a simpler data structure that most efficiently loaded each variable onto the fewest number of factors as possible. Factors with eigenvalues greater than one were included in the analysis. Chi square tests were used to determine an optimal number of factors to explain the variance. Additionally, in a subset of the F_2_ mice, we conducted a separate factor analysis in order to determine if premorbid compulsive-and anxiety-like behaviors related to BE behaviors. Because this analysis revealed no loadings between premorbid compulsive-like phenotypes and BE phenotypes, we conducted separate factor analyses for the group of premorbid versus compulsive-like variables.

### 2.8. Power analysis

We used G*Power (http://www.gpower.hhu.de/en.html) and the means, standard deviations, and unpaired two-tailed t-tests to calculate effect sizes (Cohen’s d) between homozygous B6J versus D2J genotypes, required sample sizes of homozygous genotypes based on 80% power (p < 0.05), and achieved power at each genetic locus. Because we did not have *a priori* knowledge of the mode of inheritance at each locus and because we tested a small number of F_2_ mice, we did not include heterozygous phenotypes in this analysis which yielded more conservative estimates of achieved power and required sample sizes.

## 3. RESULTS

### 3.1. BE and CPP in B6J and D2J strains

In examining strain differences in BE behaviors in a CPP paradigm (**Fig.1A**), D2J mice trained with PF (*n* = 24; 14 females, 10 males) showed greater food intake than D2J mice trained with Chow (*n* = 23; 9 females/14 males), B6J PF (*n* = 24; 12 females/12 males), and B6J Chow groups (*n* = 24; 12 females/12 males; **Fig.1B**). Both PF-trained D2J and B6J mice consumed more food than either Chow-trained group on all food days except day (D)9 for B6J (**Fig.1B**). D2J females exhibited an overall higher summed normalized food intake than males, regardless of the food type (**Fig.1C**). Examination of the slopes of escalation in normalized food intake over the course of the study revealed that the B6J PF, D2J PF, and D2J Chow groups all escalated their intake (**Fig.1D**). PF-trained D2J mice clearly exhibited the most robust escalation in intake as reflected by the largest slope (**Fig.1D**).

**Figure 1.**
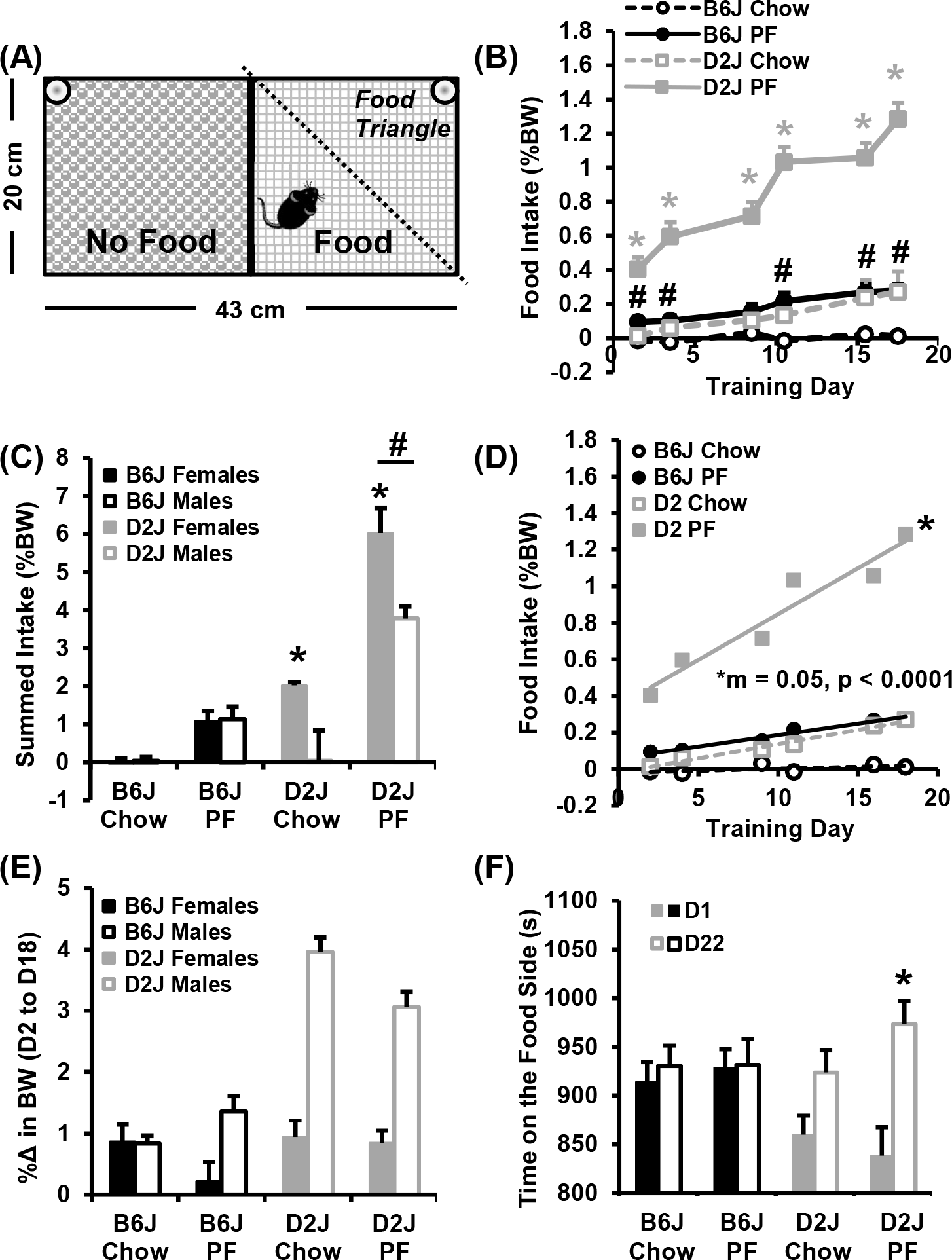
BE and CPP in B6J and D2J strains. **A)** The conditioned place preference (CPP) chamber is illustrated as seen from above with the video recording camera. **B):** A mixed effects three-way ANOVA (Genotype, Treatment, Day) revealed that D2J mice trained with PF showed significantly more food intake than all other groups [* denotes D2J PF > all other groups on all days (all ps < 0.0006); **#** B6J PF > B6J Chow on D2, D4, D11, D16, and D18 (all ps < 0.004)]. In examining PF intake, there was a main effect of Genotype [*F*_1,540_ = 269.8, *p* < 2 × 10^−16^] and Treatment [*F*_1,540_ = 321.7, *p* < 2 × 10^−16^] as well as Genotype × Day [*F*_1,540_ = 8.1, *p* = 2.2 × 10^−7^], Day × Treatment [*F*_1,540_ = 6.2, *p* = 1.2 × 10^−5^], and a Genotype × Day × Treatment interaction [*F*_1,540_ = 112.7, *p* < 2 × 10^−16^]. **C):** Females showed a higher summed normalized food intake than males overall [*F*_1,87_ = 20.4, *p* = 2.0 × 10^−5^; females > males (* *post hoc* t-test ps < 0.006); **#** D2J > all other groups]. **D):** There was an escalation in food intake in B6J PF, D2J PF, and D2J Chow groups (ps = 0.0003, 0.002, and 0.0002 vs. 0, respectively). Mice in the D2J PF group showed the most robust escalation in intake as indicated by the greatest slope (F_3,16_ = 33.3, p < 0.0001). **E):** The D2J Chow group gained more weight than all other groups (main effect of Genotype [*F*_1,87_ = 53.3, *p* = 1.29 × 10^−10^], main effect of Treatment [*F*_1,87_ = 8.3, *p* = 0.005], main effect of Sex [*F*_1,87_ = 47.8, *p* = 7.6 × 10^−10^], interaction between Genotype and Sex [*F*_1,87_ = 26.4, *p* = 1.7 × 10^−6^], interaction between Genotype, Treatment, and Sex [*F*_1,87_ = 7.1, *p* = 0.009]). **F):** Only the D2J PF group displayed an increase in time spent on the food side (right side) of the CPP box on D22 versus D1 (interaction between Genotype and Day [*F*_1,166_ = 7.5, *p* = 0.007]; *post hoc* t-test p = 0.001;* D22 > D1).

The relationship between intake of a particular diet and percent changes in body weight across Training Days was complex. Chow-trained D2J mice displayed the highest percent change in body weight compared to all other groups (Fig.1E; **Fig.S2**) yet displayed much less food intake than the D2J PF group during BE training (**Fig.1B**). Generally speaking, although PF-trained mice consumed more food during the BE training sessions, they showed less % increase in body weight than those trained with Chow. A lower percent increase in body weight could result from decreased home cage chow intake and thus, decreased caloric intake, perhaps in anticipation of the more preferred PF, as has been previously reported [19, 46, 47]. Alternatively, it could mean that there is greater energy expenditure in the home cage. Finally, although female D2J mice showed the greatest amount of food consumed for each Treatment (**Fig.1C**), they actually showed much less % increase in body weight than their male D2J counterparts (**Fig.1E**).

To examine genetic differences in conditioned food seeking behavior, we compared the time spent on the food-paired side of the CPP box during open access on D22 with the initial time spent on D1. Corresponding with the increased BE observed in PF-trained D2J mice (**Fig.1B**), these mice also displayed an increase in time spent on the food-paired side of the CPP box on D22 versus D1 (**Fig.1F**). Thus, D2J mice show both increased BE and increased conditioned reward for PF.

### 3.2. Compulsive-like eating and concomitant behaviors in B6J and D2J strains

Following assessment of conditioned food seeking behavior on D22, on D23 we assessed compulsive-like food seeking in the light/dark conflict test whereby all groups were provided access to PF located in the normally aversive light side of the light/dark box, thus creating a conflict (**Fig.2A**). PF-trained D2J mice exhibited much greater compulsive-like eating than all other groups (**Fig.2B**), consistent with an association between BE during training and compulsive-like eating. However, despite the increase in compulsive-like eating, D2J mice actually spent *less* time in the light side of the light/dark box (**Fig.2C**), and showed *fewer* entries into the light side (**Fig.2D**), as well as an increase in the number of freezing episodes (**Fig.2E**), and a shorter mean visit time in the light side (**Fig.2F**) than B6J mice. No differences in avoidance/anxiety-like behavior were found between treatment groups of each strain (**Fig.2C-F**), indicating that the increased avoidance/anxiety-like behaviors in D2J mice is driven by a stable, pre-existing genetic difference that is resistant to competing consummatory behavior. Of note, the DBA/2 strain is known to exhibit a high level of anxiety-like behavior in multiple assays, including light/dark, zero maze, and open field [48] and QTLs have been identified for anxiety-related traits in B6J × D2J-F_2_ [49] and BXD RI strains [24] indicating that strain differences in anxiety-like traits are heritable and controlled by specific genetic factors.

**Figure 2.**
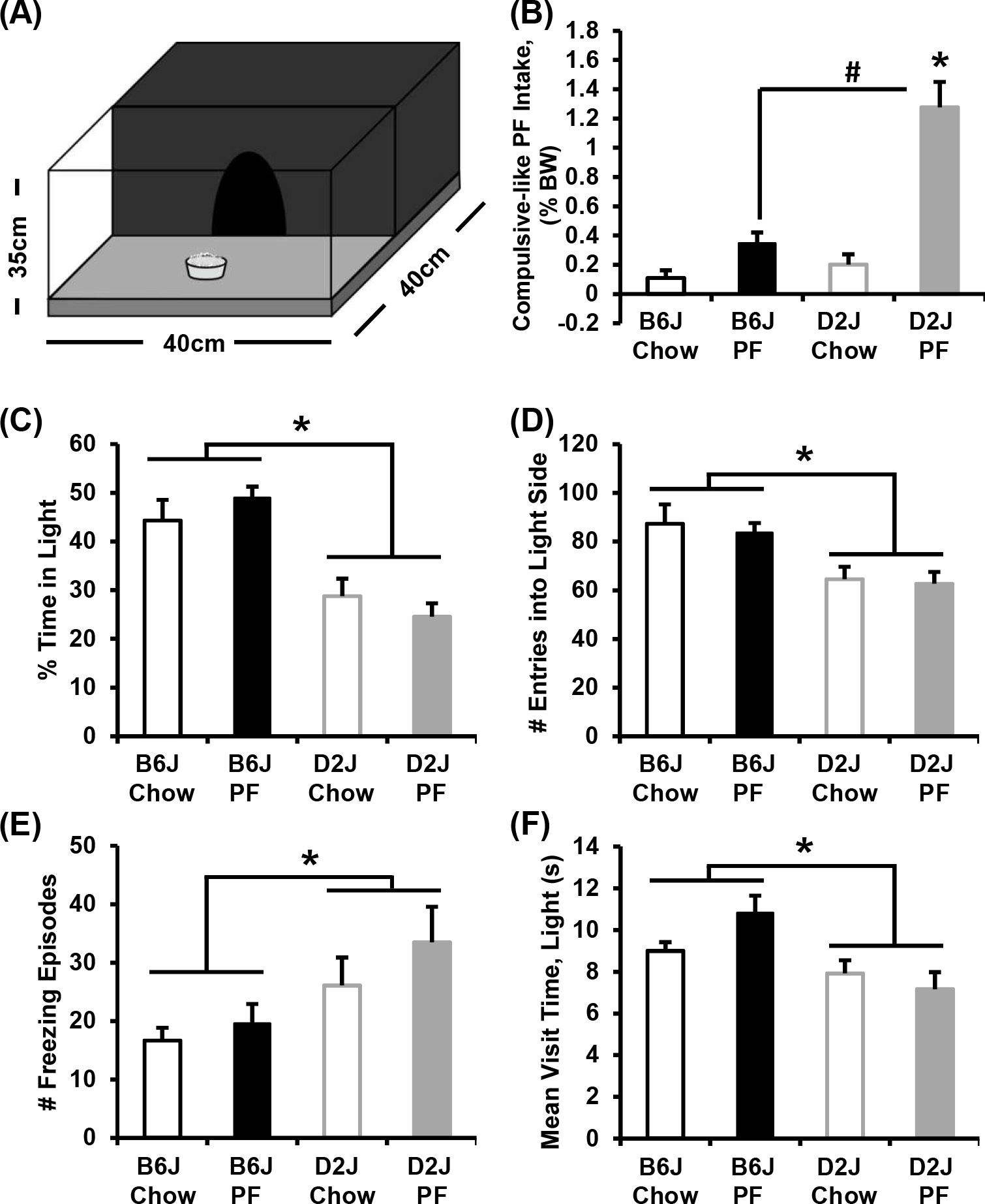
Compulsive-like eating and concomitant behaviors in B6J and D2J strains. **A):** Illustration of the light/dark conflict apparatus. **B):** A two-way ANOVA (Genotype, Treatment) found that the D2J PF group showed greater PF intake than all other groups {main effect of Genotype [*F*_1,43_ = 25.3, *p* = 9.2 × 10^−6^], main effect of Treatment [*F*_1,43_ = 40.9, *p* = 9.8 × 10^−8^], interaction between Genotype and Treatment [*F*_1,43_ = 17.6, *p* = 0.0001; *D2J PF > both Chow groups (both Bonferroni-adjusted ps < 0.00001), # D2J PF > B6J PF(Bonferroni-adjusted p < 0.00001)]. **C-F):** D2J mice also spent less time in the light side of the light/dark box (**C**; main effect of Genotype [*F*_1,43_ = 35.6, *p* = 4.1 × 10^−7^; * B6J > D2J]), showed fewer entries into the light side (**D**; main effect of Genotype [*F*_1,43_ = 14.8, *p* = 0.0004; * B6J > D2J], a greater number of freezing episodes (**E**; main effect of Genotype [*F*_1,43_ = 7.2, *p* = 0.01; * D2J > B6J], and a shorter mean visit time in the light side (**F**; main effect of Genotype [*F*_1,43_ = 11.7, *p* = 0.001; *B6J > D2J] than B6J mice.

The behavioral repertoire of compulsive-like eating in PF-trained D2J mice was in stark contrast to a recent genetic model of BE that we reported, where we found a positive, rather than negative, relationship between compulsive eating and concomitant compulsive behaviors in the BE-prone C57BL/6NJ strain [18]. Thus, PF-trained D2J mice managed to preserve their innate behavioral avoidance of light while efficiently managing to consume large amounts of PF. These results suggested a unique style of compulsive-like eating that involved a rapid entry and PF retrieval, followed by a quick retreat to the dark side where the PF was consumed. In support, D2J mice consumed a significantly greater number of pellets per visit versus B6J mice and indeed, video recording of the side view of the light side provided confirmation of an acquire-retreat-acquire-retreat strategy in D2J mice (**Fig.3** & Supplementary Information). These findings were consistent regardless of sex, in that both female and male D2J mice consumed more pellets per visit that B6J mice (data not shown).

**Figure 3.**
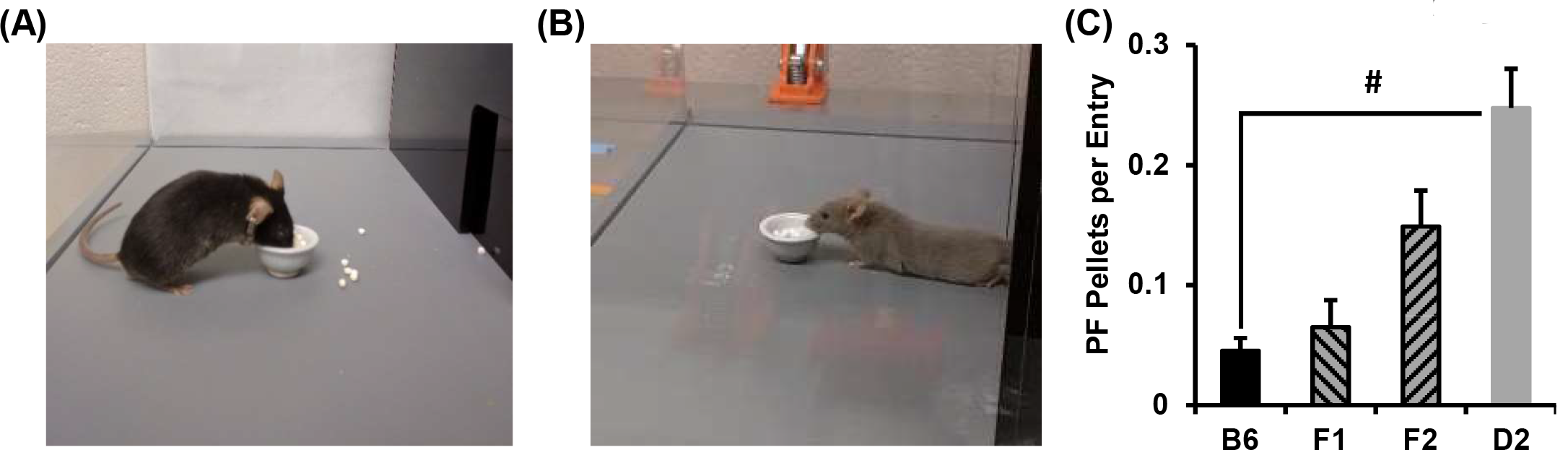
Hoarding behavior in B6J, D2J, F_1_, and F_2_ mice. **A):** B6J mice spent more time in the light and consumed their PF near the food bowl. **B)**: In contrast, D2J mice rapidly retrieved pellets and then immediately retreated to the dark side to consume them (see supplemental videos). **C):** When including the F_1_ and F_2_ data in the model, one-way ANOVA failed to detect an effect of Genotype among the groups [F_3,158_ = 0.99; p > 0.4]. When comparing the parental strains alone, D2J mice retrieved more PF pellets per entry into the light side of the Light/Dark apparatus than B6J mice, as assessed by t-test (Bonferroni-adjusted p = 3 × 10^−5^).

### 3.3. Effect sizes and power analysis

The effect size for differences in PF intake between B6J and D2J strains for Day 2 through Day 18 were quite high and increased across training days (Cohen’s d = 1.23, 1.58, 1.75, 2.28, 2.32, 2.89). In considering how many B6J × D2J-F_2_ mice to test in order to detect an effect of our three candidate loci, we assumed a much smaller medium effect size of d = 0.5 for each of our three candidate loci, resulting in a required sample size of n = 64 per homozygous genotype to achieve 80% power (p < 0.05). Assuming Mendelian ratios, n = 64 per homozygous genotype equates to generating 256 F2 mice. Because this arm of the study was a preliminary investigation into the feasibility of conducting a genome-wide QTL analysis, we tested approximately one-half of this sample size (N = 133) which provided us with 64% power to detect the effect of a locus with medium-effect size (d = 0.5; p < 0.05), assuming a Mendelian ratio and thus, a sample size yielding n = 33 per homozygous genotype.

### 3.4. Heritability estimates for BE and concomitant behaviors based on the results of the parental B6J and D2J strains

We estimated heritability (h^2^) of 16 BE-and CPP-related variables collected from PF-trained mice (**Table S1**). The most heritable phenotypes were related to PF intake. Specifically, normalized PF intake on D2, D18, and D23 in the light/dark apparatus exhibited heritability estimates of 34%, 56%, and 73%, respectively. Conversely, non-ingestive behaviors in the light/dark apparatus were the least heritable, with time in the light, number of entries into the light, and distance in the light exhibiting much lower heritability estimates ranging from 1 to 10%. Intermediately heritable phenotypes included those related to conditioned locomotor behaviors in the PF-paired side of the CPP apparatus, including D22 distance on the food side (28%), D22 entries into the food side (18%), and D22 entries into the food triangle (22%).

### 3.5. Phenotypic comparison of F_1_ and F_2_ mice with the B6J and D2J parental strains

On the first food training day, the D2J strain consumed the most PF whereasF_1_, F_2_, and B6J mice consumed a similar level (**Fig.S1A**). The D2J strain also showed greater PF intake and a greater slope of escalation than the other three groups on all days of the study. F_1_ and F_2_ mice showed slopes of escalation in PF intake that were intermediate between the D2J and B6J strains (**Fig.S1B**). These observations are consistent with a multi-locus model of overall additive inheritance, F_2_ mice showed a level of intake that was intermediate of B6J and D2J on D9, 11, 16, and 18. However, F_1_ mice showed significantly greater intake than B6J only on the final training day (D18; **Fig.S1A**). F_2_ mice also showed greater summed PF intake that B6J.F_1_ mice did not show any difference relative to B6J (**Fig.S1C**), likely due to the small sample size. D2J mice also showed greater compulsive-like PF intake than all other groups, which in turn, did not differ from each other (**Fig.S1D**). To summarize, the phenotypic effect of the B6J allele was dominant with regard to low level of initial and compulsive-like PF intake, the D2J allele dominated with regard to escalation in PF intake, and the mode of inheritance was additive with regard to intake on later training days. These observations provide evidence that different genetic factors could mediate different aspects of BE.

### 3.6. Female and male analyses of D2J versus B6J strain differences in PF intake and compulsive-like eating

We next reported strain differences separately for females and males in order to facilitate the later comparison of this data with the data showing sex-specific genetic loci. Qualitatively similar effects were observed in both sexes. In both females and males PF-trained mice showed greater intake compared to Chow-trained mice (**Fig.4A,D**). Moreover, regardless of Sex, PF-trained D2J mice showed the most robust escalation in food intake (**Fig.4B,E**), and the greatest compulsive-like PF intake (**Fig.4C,F**). Chow-trained female D2J mice showed a significant escalation in consumption whereas Chow-trained male D2J mice did not (**Fig.4B,E**), indicating less specificity of BE in female D2J mice.

**Figure 4.**
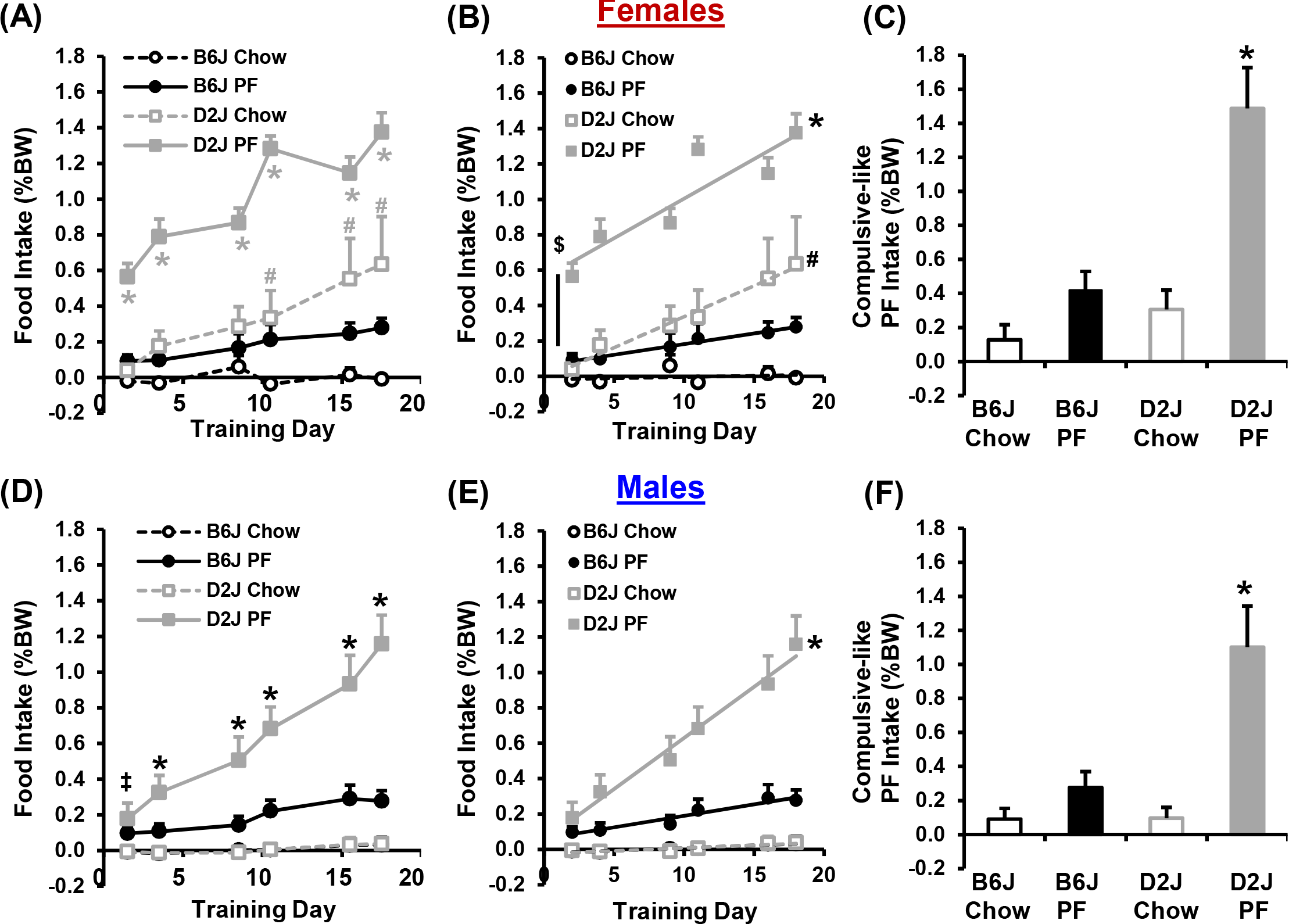
Female and male genetic differences between B6J and D2J strains in PF intake and compulsive-like eating. **A):** For female mice, there was a main effect of Genotype [F_1,258_ = 269.5, p = 2 × 10^−16^], Treatment [F_1,258_ = 169.5, p = 2 × 10^−16^], and Day [F_5,258_ = 11.5, p = 4.9 × 10^−10^], a Genotype × Treatment interaction [F_1,258_ = 41.5, p = 5.8 × 10^−10^], and a Genotype × Day interaction [F_5,258_ = 6.2, p = 2 × 10^−5^]. PF-trained D2J mice consumed more than all other groups on all days (**\***; all Bonferroni-corrected ps < 0.003), and Chow-trained D2J mice consumed more than Chow-trained B6J mice on D11, 16, and 18 (**#**; all Bonferroni-corrected ps < 0.03). **B):** PF-trained female D2J mice showed a greater slope than either of the female B6J groups (both ps <0.0001; * vs. Chow-and PF-trained B6J groups), and a greater y-intercept than the Chow-trained D2J group (**$**; p < 0.0001). Furthermore, Chow-trained female D2J mice showed a greater slope than Chow-trained B6J mice (**#**; p = 0.001). **C):** When considering compulsive-like consumption in the light/dark conflict test, there was a main effect of Treatment [F_1,19_ = 24.1; p = 9.7 × 10^−5^], Genotype [F_1,19_ = 18.4; p = 0.0004], and an interaction [F_1,19_ = 10.0; p = 0.005]. *Post hoc* pair-wise comparisons revealed greater compulsive-like consumption for the PF-trained D2J mice compared to all other groups (**\***; all Bonferroni-corrected ps < 0.0003). **D):** For male mice, there was a main effect of Genotype [F_1,264_ = 218.2; p = 2 × 10^−16^], Treatment [F_1,264_ = 60.9; p = 1.4 × 10^−13^], and Day [F_5,264_ = 11.9; p = 2.3 × 10^−10^], and a Genotype × Treatment interaction [F_1,264_ = 17.8; p = 1.7 × 10^−15^], a Treatment × Day interaction [F_5,264_ = 9.3; p = 3.4 × 10^−8^], a Genotype × Day interaction [F_5,264_ = 4.4; p = 0.0007], and a Genotype × Treatment × Day interaction [F_5,264_ = 5.1; p = 0.0002]. PF-trained male D2J mice consumed more food on D2 than either Chow-trained male group (‡; both Bonferroni-corrected ps < 0.02), and more food than all three other groups on D4-18 (*; all Bonferroni-corrected ps < 0.02). **E):** PF-trained D2J male mice showed a significantly greater slope than all other groups (*; all ps < 0.0001). **F):** In examining compulsive-like intake in male mice, there was a main effect of Treatment [F_1,20_ = 18.7; p = 0.0003], Genotype [F_1,20_ = 9.1; p = 0.007], and an interaction [F_1,20_ = 8.9; p = 0.007]. PF-trained D2J male mice showed greater compulsive-like consumption than any of the other three groups (*; all corrected ps < 0.002).

### 3.7. Genotypic analysis of candidate loci on chrs. 4, 6, and 11 in BE and compulsive-like eating

To provide initial insight into the genetic basis underlying the parental B6J and D2J strain differences in BE, we tested a small cohort of B6J × D2J F_2_ mice. Mice were genotyped at the peak locations of three different QTLs (see Methods) previously associated with sweet taste (chr. 4; rs6316711, 156 Mb), bitter taste (chr. 6; rs13479039; 133 Mb), and sensitivity to drugs of abuse (chr. 11; rs 29383600; 50 Mb). For the sweet taste locus on chromosome 4, we did not identify any significant effect of Genotype on PF intake over time (**Fig.5A**), slope of escalation of intake (**Fig.5B**), or compulsive-like PF intake in the light/dark apparatus (**Fig.5C**).

**Figure 5.**
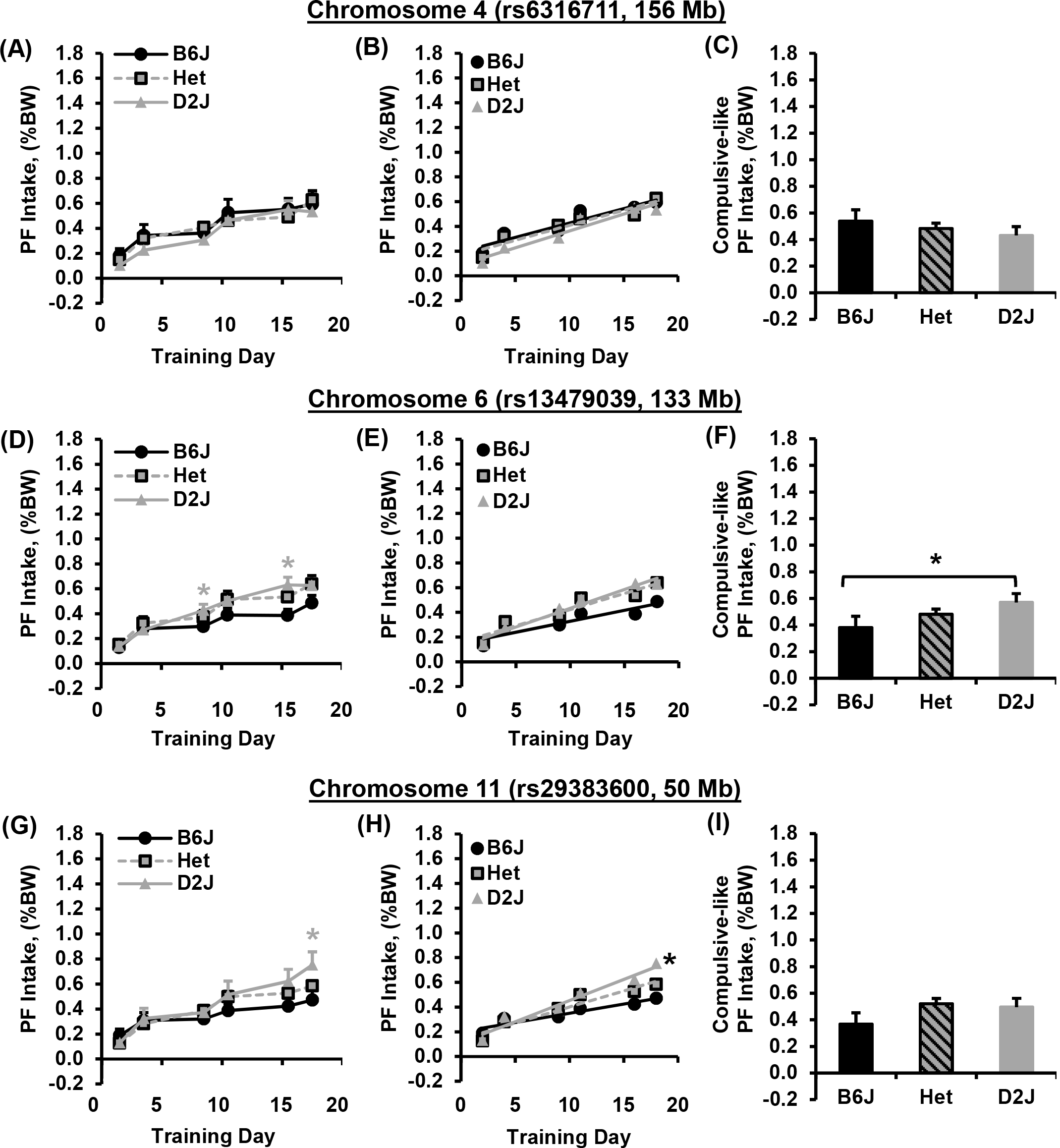
Effect of Genotype at three candidate loci on BE and compulsive-like eating in B6J × D2J-F_2_ mice. **A-C):** For the chromosome 4 locus (sweet taste), there was no effect of Genotype on PF intake (**A**; p = 0.16), escalation slope (**B**; p = 0.78), or compulsive-like PF intake in the light side of the light/dark box (**C**; p = 0.52). **D-F):** For the chromosome 6 locus (bitter taste), there was an effect of Genotype [**D**; F_2,780_ = 6.4, **\***p = 0.002] in that mice homozygous for D2J consumed more PF than mice homozygous for B6J mice on D9 and D16 (p = 0.04 and 0.003, respectively). There was no effect of Genotype on slope of escalation [**E**; F_2,786_ = 2.5, p = 0.08] and all three genotypes showed a significant, non-zero slope (all p < 0.0001). For compulsive-like PF intake, of the effect of Genotype was not significant [**F**; F_2,129_ = 2.5, p = 0.08]; however, mice homozygous for D2J consumed more PF than mice homozygous for B6J as assessed via t-test (*p = 0.02). **G-I):** For the chromosome 11 locus (behavioral sensitivity to drugs of abuse), there was an effect of Genotype on PF intake [**G**; F_2,780_ = 3.7, p = 0.02] in that mice homozygous for D2J showed greater intake than mice homozygous for B6J on D18 (**\***p = 0.03). Additionally, mice homozygous for D2J showed a greater slope of escalation in PF intake compared to mice homozygous for B6J [**H**; *; F_2,786_ = 4.3, p = 0.01]. For compulsive-like intake, there was no effect of Genotype (**I**; p = 0.15).

For the bitter taste locus on chromosome 6, mice homozygous for the D2J allele showed a significant increase in PF intake on D9 and D16 relative to mice homozygous for the B6J allele (D2J vs. B6J; **Fig.5D)**. The effect sizes between the two homozygous genotypes were Cohen’s d = 0.48 and 0.7, respectively and with n = 38 B6J and n = 39 D2J, we achieved 55% and 86% power, respectively. The chromosome 6 locus explained 5% and 11% of the phenotypic variance in PF intake on D9 and D16. The h^2^ of D9 and D16 phenotypes was 43% and 57%, respectively. The chromosome 6 locus explained 5% and 10% of the genetic variance in PF intake on D9 and D16. (**Table 1**). There was no effect of Genotype on slope of escalation (**Fig.5E**). For compulsive-like intake, mice homozygous for D2J consumed more PF than mice homozygous for B6J (**Fig.5F**). The h^2^ of compulsive-like eating was 69% and the chromosome 6 locus explained 7% of the phenotypic variance and 3% of the genetic variance in compulsive-like eating (**Table 1**).

**Table 1.**
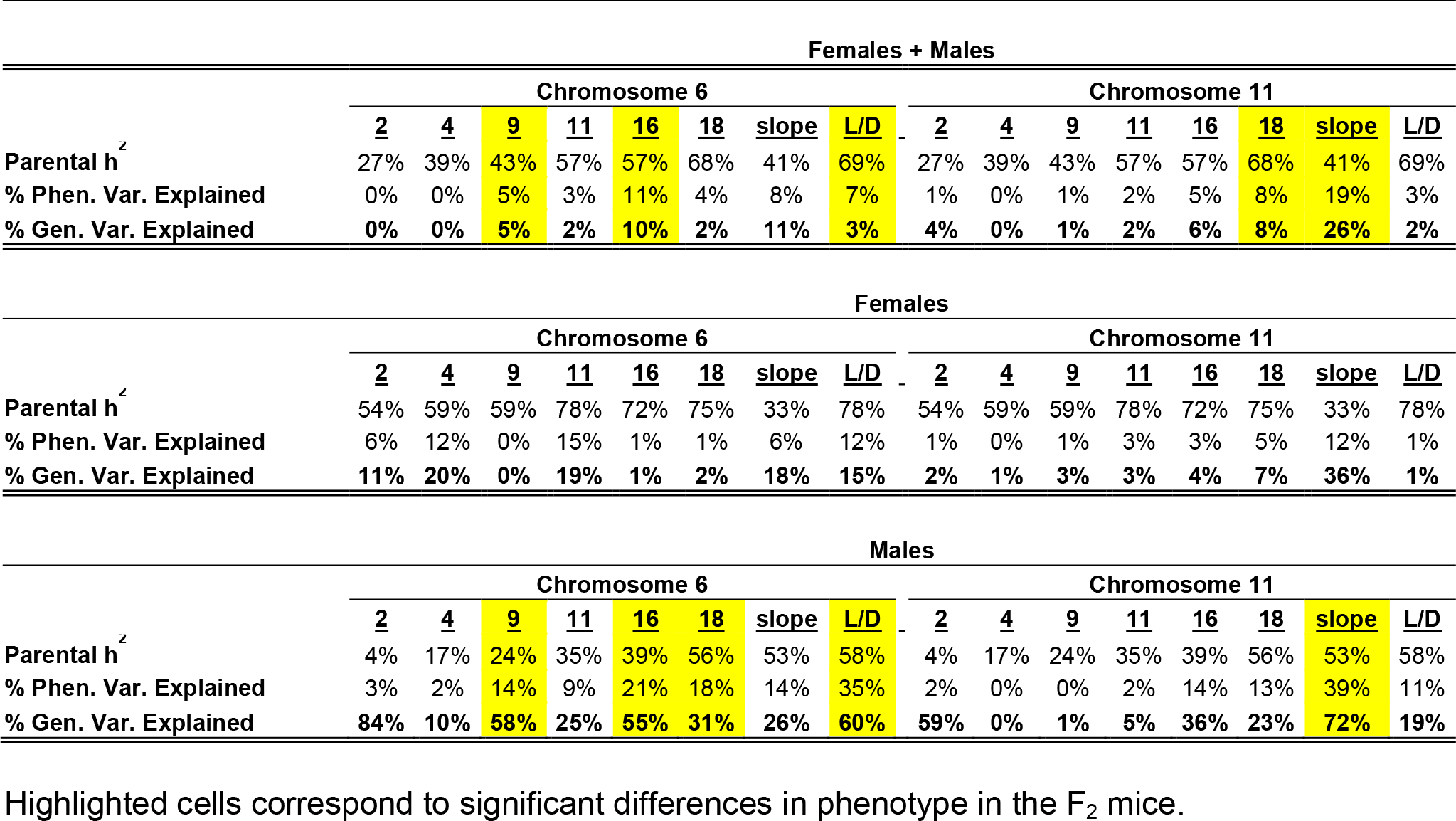
Heritability estimates and genetic variance in B6/D2-F_2_ mice.

| Females + Males |  |  |  |  |  |  |  |  |  |  |  |  |  |  |  |  |
| --- | --- | --- | --- | --- | --- | --- | --- | --- | --- | --- | --- | --- | --- | --- | --- | --- |
|  | Chromosome 6 |  |  |  |  |  |  |  | Chromosome 11 |  |  |  |  |  |  |  |
|  | <u>2</u> | <u>4</u> | <u>9</u> | <u>11</u> | <u>16</u> | <u>18</u> | <u>slope</u> | <u>L/D</u> | <u>2</u> | <u>4</u> | <u>9</u> | <u>11</u> | <u>16</u> | <u>18</u> | <u>slope</u> | <u>L/D</u> |
| Parental h<sup>2</sup> | 27% | 39% | 43% | 57% | 57% | 68% | 41% | 69% | 27% | 39% | 43% | 57% | 57% | 68% | 41% | 69% |
| % Phen. Var. Explained | 0% | 0% | 5% | 3% | 11% | 4% | 8% | 7% | 1% | 0% | 1% | 2% | 5% | 8% | 19% | 3% |
| % Gen. Var. Explained | 0% | 0% | 5% | 2% | 10% | 2% | 11% | 3% | 4% | 0% | 1% | 2% | 6% | 8% | 26% | 2% |
| Females |  |  |  |  |  |  |  |  |  |  |  |  |  |  |  |  |
|  | Chromosome 6 |  |  |  |  |  |  |  | Chromosome 11 |  |  |  |  |  |  |  |
|  | <u>2</u> | <u>4</u> | <u>9</u> | <u>11</u> | <u>16</u> | <u>18</u> | <u>slope</u> | <u>L/D</u> | <u>2</u> | <u>4</u> | <u>9</u> | <u>11</u> | <u>16</u> | <u>18</u> | <u>slope</u> | <u>L/D</u> |
| Parental h<sup>2</sup> | 54% | 59% | 59% | 78% | 72% | 75% | 33% | 78% | 54% | 59% | 59% | 78% | 72% | 75% | 33% | 78% |
| % Phen. Var. Explained | 6% | 12% | 0% | 15% | 1% | 1% | 6% | 12% | 1% | 0% | 1% | 3% | 3% | 5% | 12% | 1% |
| % Gen. Var. Explained | 11% | 20% | 0% | 19% | 1% | 2% | 18% | 15% | 2% | 1% | 3% | 3% | 4% | 7% | 36% | 1% |
| Males |  |  |  |  |  |  |  |  |  |  |  |  |  |  |  |  |
|  | Chromosome 6 |  |  |  |  |  |  |  | Chromosome 11 |  |  |  |  |  |  |  |
|  | <u>2</u> | <u>4</u> | <u>9</u> | <u>11</u> | <u>16</u> | <u>18</u> | <u>slope</u> | <u>L/D</u> | <u>2</u> | <u>4</u> | <u>9</u> | <u>11</u> | <u>16</u> | <u>18</u> | <u>slope</u> | <u>L/D</u> |
| Parental h<sup>2</sup> | 4% | 17% | 24% | 35% | 39% | 56% | 53% | 58% | 4% | 17% | 24% | 35% | 39% | 56% | 53% | 58% |
| % Phen. Var. Explained | 3% | 2% | 14% | 9% | 21% | 18% | 14% | 35% | 2% | 0% | 0% | 2% | 14% | 13% | 39% | 11% |
| % Gen. Var. Explained | 84% | 10% | 58% | 25% | 55% | 31% | 26% | 60% | 59% | 0% | 1% | 5% | 36% | 23% | 72% | 19% |

Highlighted cells correspond to significant differences in phenotype in the F<sub>2</sub> mice.

For the locus influencing sensitivity to drugs of abuse on chromosome 11, mice homozygous for D2J consumed more PF than mice homozygous for B6J on D18 (**Fig.5G**). The effect size of D18 intake was d = 0.58 and with n = 32 homozygous B6J and n = 27 homozygous D2J samples, we achieved 59% power. The h^2^ of D18 intake was 68% and the chromosome 11 locus (50 Mb) explained 8% of the phenotypic variance and 8% of the genetic variance in D18 PF intake (**Table 1**). Mice homozygous for D2J also showed a greater slope of escalation in PF intake compared to mice homozygous for B6J (**Fig.5H**), but no difference in compulsive-like PF intake (**Fig.5I**). The h^2^ of the slope of PF intake was 41% and the chromosome 11 locus explained 19% of the phenotypic variance and 26% of the genetic variance (**Table 1**).

### 3.8. Male-specific effect of the bitter taste locus on chromosome 6

When we considered the effect of Genotype at chromosome 6 separately in each sex, for females, there was no effect of Genotype on PF intake (**Fig.6A-C**). In contrast, males completely accounted for the observed effect in which homozygous for the D2J allele showed greater PF intake on D9, D16, and D18 (**Fig.6D**), a greater slope of escalation (**Fig.6E**), and a greater compulsive-like PF intake (**Fig.6F**) relative to males homozygous for the B6J allele. The h^2^ of the significant male-specific phenotypes ‑ D9, D16, D18, and compulsive-like consumption ‑ was 24%, 39%, 57%, and 58% (**Table 1**). The percent phenotypic variance explained at the chromosome 6 locus for these four male-specific phenotypes was 14%, 21%, 18%, and 35% and the percent genetic variance explained was 58%, 55%, 31%, and 60% (**Table 1**). Thus, the chromosome 6 locus is a major, male-specific locus influencing BE.

**Figure 6.**
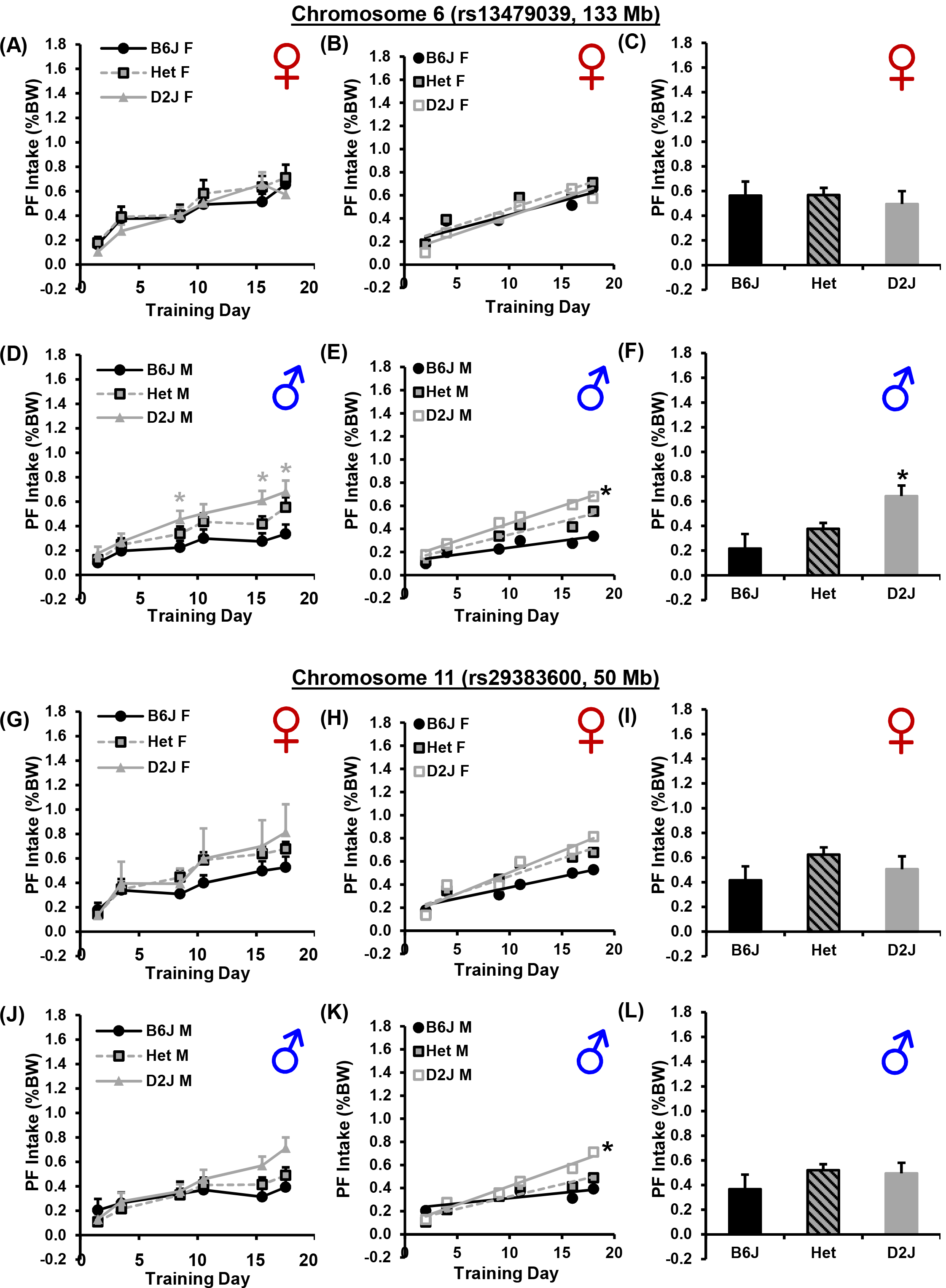
Effect of Genotype at three candidate loci on BE and compulsive-like eating in female and male B6J × D2J-F_2_ mice. **A):** For females at the chromosome 6 locus (bitter taste), there was no effect of Genotype and no interaction with Day (both ps > 0.3). **B):** There were no genotypic differences in the slope of escalation [F_2,384_ = 0.3; p = 0.76]. All female genotypes showed significant slopes (all ps < 0.0001 vs. 0). **C):** There were no genotypic differences in compulsive-like PF consumption (p > 0.7). **D):** For males at the chromosome 6 locus (bitter taste), there was a main effect of Genotype [F_2,372_ = 14.0; p = 1.3 × 10^−6^] but no interaction with Day (p > 0.6). Males homozygous for D2J consumed more PF than males homozygous for B6J on D9, D16, and D18 (*; all Bonferroni-corrected ps < 0.05). **E):** There was a significant slope in escalation for all three male genotypes (all ps < 0.008 vs. zero) and greater escalation in males homozygous for D2J compared to males homozygous for B6J (**\***; Bonferroni-corrected p = 0.02). **F):** For compulsive-like PF consumption in males, there was an effect of Genotype [F_2,62_ = 9.6; p = 0.0002], in that males homozygous for D2J consumed more than the other genotypes (*; both Bonferroni-corrected ps < 0.04). **G):** For females at the chromosome 11 locus, there was a main effect of Genotype [F_2,372_ = 3.1; p = 0.047]. However, when we compared PF consumption on each day, the effect of Genotype did not withstand adjustment for multiple comparisons (all Bonferroni-adjusted ps > 0.2). **H):** There was no genotypic difference in the slope of escalation among females [F_2,396_ = 1.7; p = 0.18). **I):** There was no genotypic difference in compulsive-like PF consumption among females (p > 0.14). **J):** For males at the chromosome 11 locus, there was an effect of Genotype [F_2,372_ = 3.3; p = 0.04]. Similar to observations in female mice, this effect did not withstand multiple comparisons on individual days (all ps > 0.1). **K):** There was a significant slope in males homozygous for D2J and in heterozygous males (both ps < 0.0001 vs. zero), but not in males homozygous for B6J (p > 0.19). D2J males showed a significantly greater slope than B6J males (*; Bonferroni-adjusted p = 0.02). **L):** There was no difference in compulsive-like PF intake among the male genotypes (p > 0.3).

### 3.9. Chromosome 11 locus: females versus males

For females, the effect of Genotype on PF intake at the chromosome 11 locus was not significant during training or compulsive-like assessment (**Fig.6G-I**). For males, mice homozygous for the D2J allele showed a significant increase in slope of escalation of PF intake relative to mice homozygous for the B6J allele (**Fig.6K,L**), thus accounting for the overall effect of Genotype in **Fig.5H**. The h^2^ of this trait was 53% in males and the chromosome 11 locus explained 39% of the phenotypic variance and 73% of the genetic variance in the slope of escalation in PF intake (**Table 1**).

### 3.10. Exploratory factor analysis of BE and compulsive eating in B6J, D2J, and F_2_ mice

To reveal potential novel relationships among the 16 major BE-and CPP-related variables, we conducted exploratory factor analysis in PF-trained B6J, D2J, and F_2_ mice (**Table S1**). The top three factors for each group are color coded according to shared variables for each factor (not according to percent variance explained). Here, we discuss variables with factor loadings of 0.4 or greater. For the B6J strain (**Fig.7A**), three factors explained 55% of the total variance [χ^2^ (88, *N* = 24) = 158.8; p = 5.7 × 10^−6^]. For the D2J strain (**Fig.7B**), three factors explained 57% of the variance [χ^2^ (88, *N* = 24) = 155.2; p = 1.3 × 10^−5^].

**Figure 7.**
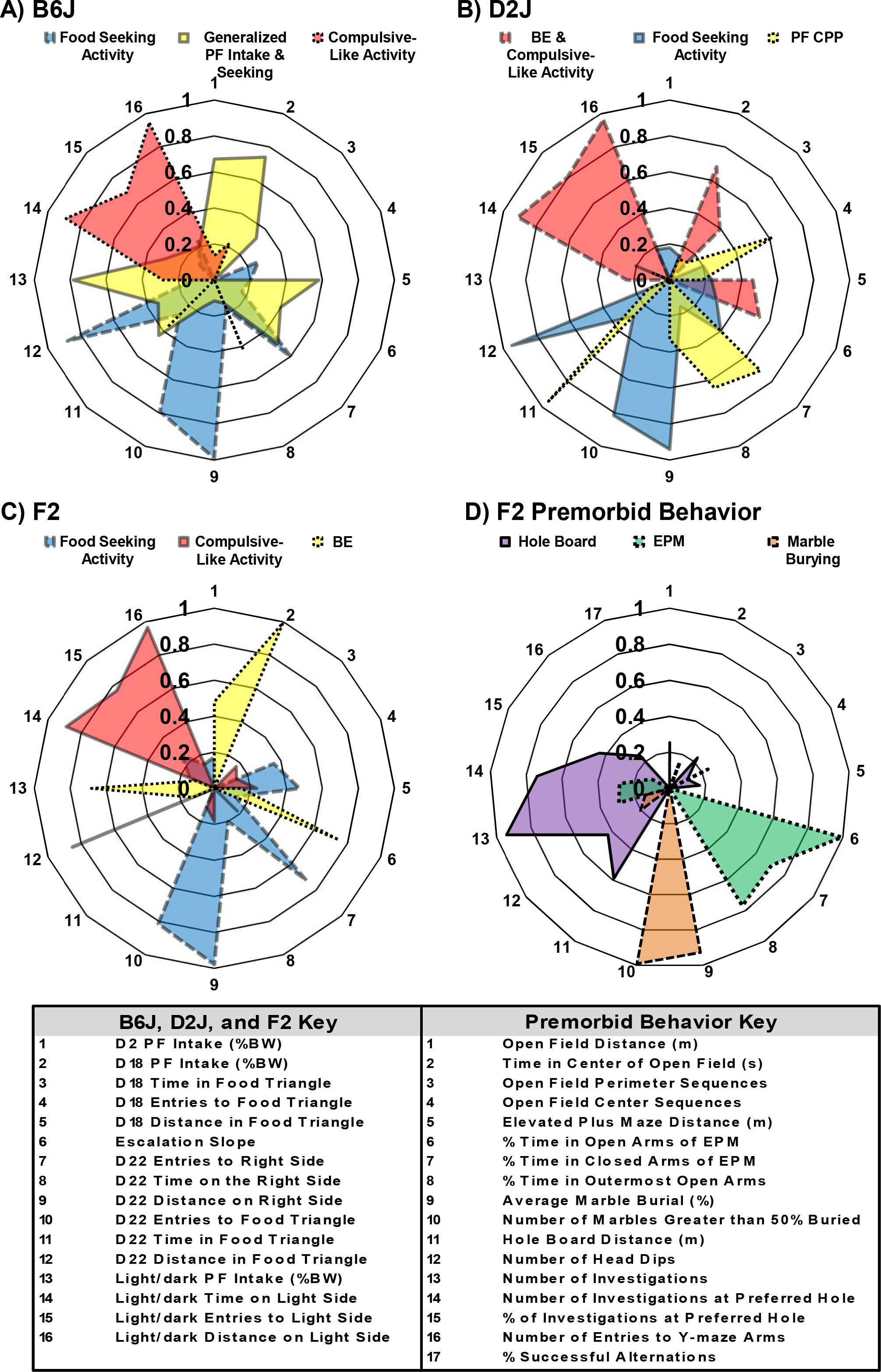
Factor analysis of BE-associated phenotypes in B6J, D2J, and B6J × D2J-F_2_ mice. Data are illustrated as the absolute value of loadings for each variable for ease of presentation. For additional details, including whether the loadings were positive or negative, see Table S1. **A):** For the B6J inbred strain, the blue factor explained 19% of the variance and comprised phenotypes related to CPP, namely D22 Entries to the Food Side, D22 Distance on the Food Side, D22 Entries to the Food Triangle, and D22 Distance in the Food Triangle. This factor was termed “Food Seeking Activity”. The yellow factor (Generalized PF Seeking & Intake) explained 19% of the variance related to PF seeking and intake and encompassed the variables D2, D18, and Light/Dark PF intake. The red factor (Compulsive-like Activity) explained 17% of the variance in compulsive-related behaviors and encompassed Time on, Entries to, and Distance on the Light Side. **B):** For the D2J inbred strain, the red factor (BE & Compulsive-like Activity) explained 23% of the variance related to BE and compulsive behaviors and encompassed D18 PF Intake, Escalation Slope, and three non-ingestive compulsive behaviors in the Light/Dark apparatus. The blue factor (Food Seeking Activity) explained 19% of the variance and comprised CPP-related variables, including D22 Distance on the Right Side and in the Food Triangle as well as Entries to the Food Triangle. The yellow factor (PF CPP) explained 15% of the variance and comprised activity variables related to PF seeking, including D18 Entries to the Food Triangle, D22 Entries to the Right Side, and D22 Time in the Food Triangle. **C):** For F_2_ mice, the blue factor (Food Seeking Activity) explained 21% of the variance and encompassed factors related to Conditioned Place Preference, namely D22 Entries to the Right Side, D22 Distance on the Right Side, D22 Entries to the Food Triangle, and D22 Distance in the Food Triangle. The red factor (Compulsive-Like Activity) explained 16% of the variance encompassing variables related to compulsive behavior, specifically Time on, Entries into, and Distance on the Light Side of the Light/Dark apparatus. The yellow factor (Generalized Intake, BE, and Compulsive-Like BE) explained 15% of the variance related to BE and included D2 and D18 PF Intake, as well as PF Intake in the Light/Dark apparatus. **D):** For premorbid compulsive-like behavior in F_2_ mice, the purple factor explained 14% of variance related to measures in the hole board test. The green factor explained 14% of the variance and encompassed three of the four EPM measures, including % Time in the Open Arms, % Time in the Closed Arms, and % Time in the Outermost Area of the Open Arms. The brown factor explained 11% of the variance and encompassed both measures of the marble burying test.

The blue factor explained 19% of the variance in both B6J mice and D2J mice. We termed this factor “Food Seeking Activity” for both strains, as it encompassed four variables related to activity during CPP assessment on D22 (#7, #9, #10, and #12), including D22 Entries to the Food Side (#7), D22 Distance on the Food Side (#9), D22 Entries to the Food Triangle (#10), and D22 Distance in the Food Triangle (#12).

The red factor in the B6J strain explained 17% of the variance and in D2J explained 23% of the variance and comprised an overlapping yet largely distinct set of variables between the two strains. For B6J mice, the red factor only comprised the three main compulsive behavior variables (#14, #15, and #16), including Light/Dark Time (#14), Light/Dark Entries (#15), an Light/Dark Distance (#16) and was termed “Compulsive-Like Activity.” For the D2J strain, in addition to these three variables, the red factor also contained D18 PF Intake (#2), D18 Time in Food Triangle (#3), D18 Distance in Food Triangle (#5), and Escalation Slope (#6) and for this reason, was termed “BE and Compulsive-Like Activity”.

The yellow factor in the B6J strain explained 19% of the variance in the D2J strain explained 15% of the variance and was also largely distinct between the two strains. For the B6J strain, the yellow factor was more extensive and contained 7 variables with loadings greater than 0.4 (#1, #2, #5, #6, #7, #11, and #13), including D2 PF Intake (#1), D18 PF Intake (#2), D18 Distance in Food Triangle (#5), Escalation Slope (#6), D22 Entries to the Food Side (#7), D22 Time in Food Triangle (#11), Light/Dark PF Intake (#13) and was termed “Generalized PF Intake and Seeking.” The generalized intake refers to the fact that acute D2 intake (prior to binge escalation) loaded onto a common intake factor in B6J mice.

The yellow factor in D2J mice accounted for less variance and only contained 4 variables with loadings greater than 0.4 (#4, #7, #8, #11) that also included D22 Entries to the Food Side (#7) and D22 Time in Food Triangle (#11) but also D18 Entries into the Food Triangle (#4) and D22 Time on the Food Side (#8). However, the yellow factor in D2J mice did not include any intake variables (those variables loaded onto the red factor) and thus, was termed “PF CPP”. To summarize, the most striking finding from exploratory factor analysis of the parental strains was that for B6J mice, intake and concomitant activity variables loaded onto a single factor whereas in D2J mice, only some of the intake variables loaded onto a single factor which also contained subsequent compulsive-like behaviors.

We next conducted exploratory factor analysis in 133 F_2_ mice that segregated B6J and D2J alleles. The purpose of this analysis was to determine the extent to which variation among behavioral variables was co-inherited as evidenced by co-loading of variables onto a common factor versus independent inheritance as evidence by distinct loading onto separate factors. In F_2_ mice, three factors explained 52% of the total variance [χ^2^ (75, *N* = 133) = 456.4; p = 5.0 × 10^−56^; **Fig.7C**].

The blue factor (#5, #7, #9, #10, and #12) in F_2_ mice accounted for 21% of the total variance and was similarly termed “Food Seeking Activity”, as it encompassed largely the same phenotypes as the parental strains (in particular B6J) related to activity in the food-paired side and the food triangle on the preference assessment day (D22) and additionally, D18 Distance in the Food Triangle.

The red factor (#14, #15, and #16) in F_2_ mice explained 16% of the variance and similar to the B6J strain was termed “Compulsive-Like Activity” as it encompassed the same activity measures in the light/dark apparatus.

The yellow factor (1,2,6,13) explained 15% of the variance, and was clearly distinct from two parental strains as it purely contained all of the intake phenotypes, including D2 Intake, D18 Intake, Escalation Slope, and Light/Dark Intake (D23). This factor was therefore termed “BE”.

### 3.11. Exploratory factor analysis of premorbid compulsive/anxiety-like battery in F_2_ mice

We also analyzed premorbid compulsive and anxiety-like behaviors separately in a subset of the F_2_ mice. Three factors explained 40% of the total variance [χ^2^ (88, *N* = 93) = 169.7; p = 3.9 × 10^−7^; **Fig.7D**]. The purple factor explained 14% of the variance and consisted of behaviors in the hole board test. The green factor explained 14% of the variance, encompassed behaviors in the EPM test. The brown factor explained 11% of the variance and contained the two marble burying phenotypes. None of the open field variables or y-maze variables loaded onto any factor. Together, these results indicate that there is little overlap variance explained between the behavioral variables of the five assays comprising the anxiety/compulsive-like battery and suggest that the genetic basis of each assay is likely to be largely distinct. Furthermore, the lack of co-loading of any of these premorbid variables onto the BE ingestive and non-ingestive measures suggest that they are not genetically correlated with BE and thus, lack predictive value in this F_2_ cross.

## 4. DISCUSSION

We identified robust strain differences in BE, with the D2J strain but not the B6J strain exhibiting a dramatic escalation in PF consumption along with compulsive-like eating in the light-dark test (**Figs.1-3**). Consistent with our previous studies of BE in B6J mice [18] or WT littermates on a B6J background [25], B6J mice showed little evidence of BE in our intermittent, limited access paradigm of sweetened PF and correspondingly, showed no conditioned PF reward or compulsive-like eating (**Figs.1,2**). In examining the behavioral organization of BE and other behavioral measures in the parental strains, for the BE-resistant B6J strain, ingestive behaviors (yellow: #1, #2, #6) loaded onto a separate factor from light/dark activity (red; #14, #15, #16; **Fig.7A**). In contrast, for the BE-prone D2J strain, neither initial PF intake (#1) nor compulsive-like eating (#13) loaded onto any factor whereas BE-level intake (D18) and slope of escalation loaded onto the same factor as compulsive-like activity – a factor that we termed “BE and Compulsive-Like Activity” for the D2J strain (red; **Fig.7B**). Together, these findings that the unique hoarding-like behavior in the light/dark box in the D2J strain (**Fig.3**) was reflective of the degree of BE (but not initial PF ingestion), whereas in B6J mice, simple initial intake was more predictive of future intake. In contrast to the parental strains, factor analysis in the F_2_ mice indicated that all PF intake variables, including intake on D2 and D18, slope of escalation, and compulsive-like intake, loaded cleanly onto a single factor (yellow; **Fig.7C**). None of the premorbid anxiety/compulsive-like factors loaded onto any of the PF consumption phenotypes in the initial analysis (not shown) nor did they overlap in their factor loadings (**Fig.7D**), suggesting a separate genetic basis from BE as well as a separate genetic basis for each of these premorbid assays.

Strain differences in BE could not be explained by initial differences in body weight as there was no difference on the first day of the training session **(Fig.S2)**, yet there was already a marked increase in PF consumption in D2J mice (**Fig.1B; Fig.4A,D; Fig.S1A**). Furthermore, the D2J increase in PF intake was accompanied by *reduced* weight gain relative to Chow-trained D2J mice who gained significantly more weight (PF = 7.6% ± 1.1%; Chow = 11.6% ± 1.4%, t(45) = 2.2; p = 0.03; see **Fig.S2**). The observation that BE was associated with *less* percent change in BW is further supported by D2J females showing a 1.5-fold greater increase in % BW consumed compared to D2J males (**Fig.1C**), yet showing a three-fold lower percent increase in % BW relative to male mice (**Fig.1E**). To summarize, differences in body weight are not associated with the initial difference in PF consumption and the escalation in consumption is associated with a reduced % increase in BW. The latter conclusion is further supported by a negative correlation between the total % BW consumed and change in BW in F_2_ mice (r = −0.24; p = 0.006)

It is interesting to note that despite the D2J strain exhibiting a large degree of compulsive-like consumption of PF, they nevertheless continued to exhibit marked avoidance/aversion for the light side of the light/dark box, showing less time, fewer entries, and a shorter average visit time that was similar to Chow-trained D2J mice (**Fig.3**). This is in stark contrast to the BE-prone B6NJ strain (a closely related substrain of B6J) that exhibited an increase in compulsive-like eating, a decrease in avoidance behavior (increased time in the light side), an increase in visit time in the light side, and a concomitant fewer number of entries into the light side [18]. Clearly, the innate avoidance behavior in the D2J strain was not compromised by the profound, rapid consumption of the available PF in the open arena. In other words, the D2J strain managed to avoid the light side while at the same time, efficiently transport large amounts of PF to the safer environment where they consumed it. In support, horizontal video observation of a subset of the D2J strain confirmed that when D2J mice entered the light side, they exhibited a stretched posture and efficiently traveled as little distance as possible directly toward the food bowl, retrieved the PF, and immediately retreated back to the dark side to consume it (**Fig.3A,B**; see also Supplementary Information for videos). Furthermore, the D2J strain showed a five-fold increase in the number of pellets retrieved per entry into the light (**Fig.3C**). This behavior is in some ways reminiscent of hoarding behavior exhibited by many patients with BE [6, 50], including hoarding of food [51] with more severe hoarding behavior being associated with the presence of ED, in particular with females [52]. Hoarding may represent a behavioral strategy in addition to BE that could serve to alleviate negative emotions [50]. In humans, hoarding of food for BE is an adaptive behavior that serves to avoid the aversive social stigma associated with BE [53]. In the D2J strain, hoarding-like behavior was also adaptive since it permitted efficient intake of the PF that was on par with the amount exhibited during escalated consumption (**Fig.3B vs. Fig.1B**) without sacrificing any avoidance behavior of the aversive (and potentially dangerous) light environment. That escalated PF intake did not compete with innate avoidance behavior in PF-versus Chow-trained D2J mice is remarkable and illustrates the extreme motivation to obtain the PF and retreat as quickly as possible.

We are not aware of any other preclinical, forward genetic studies of BE besides our recent study [18]. However, there is an extensive literature on forward genetic studies of consumption of and preference for PF (e.g., sucrose and fat) in rodents and other animal species [54]. Variation in the hedonic score for sucrose shows a heritability of approximately 50% in adult humans [55], indicating a strong genetic component. Because we utilized the 5-TUL diet in which sucrose is the primary palatable ingredient, a discussion is warranted on the literature involving differences in sucrose consumption between B6J and D2J alleles [26–31, 56]. An important observation is that the length of access to sucrose can determine the directionality of strain difference in liquid sucrose consumption between B6J and D2J. Similar to our results, short-term access (2h or less) induced greater consumption in D2J [27] and longer-term access (6h or greater) induced greater consumption in B6J [26, 56, 57]. For the long-term access (B6J strain > D2J strain; opposite of our results), a QTL on distal chromosome 4 was identified [26] that was hypothesized to be mediated by one or more polymorphisms in the *Tas13r* gene [31, 56, 58]. However, there was no genetic correlation between short-and long-term consumption among inbred strains [27], suggesting that short-and long-term consumption are mediated by separate genetic factors.

We tested the effect of Genotype at the chromosome 4 *Tas13r* sweet taste locus on PF consumption in 133 B6J × D2J-F_2_ mice and were surprised to find no effect of Genotype at any time point in the BE training session (**Fig.5**) *TAS1R3* codes for one of three receptors of the *TAS1R* gene family (*TAS1R1*-*TAS1R3*). *TAS1R3* dimerizes with either *TAS1R1* or *TAS1R2* to form the sweet receptor or the umami receptor, respectively [59]. Our negative results suggested that in our paradigm, polymorphisms at the *Tas13r* locus do not contribute (at least, not additively) to strain differences in acute or escalated PF intake of the 5-TUL diet in B6J versus D2J strains. It is possible that under different conditions (e.g., longer access to the 5-TUL diet) that this locus could affect acute or escalated PF intake. Alternatively, the locus could interact epistatically with other loci and we are not powered to detect epistatic interactions with our small sample size.

Unexpectedly, we found evidence that the second taste QTL on chromosome 6, the *Tas2r* locus that was previously associated with greater consumption of a bitter solution with the D2J allele [26], was associated with greater PF intake over time and compulsive-like intake in the light-dark test (**Fig. 5D-F)**. The increased PF intake associated with the D2J allele was driven exclusively by males (D9, D16, D18, and compulsive-like eating; **Fig.6D-F**) and the *Tas2r* locus accounted for 14% to 38% of the phenotypic variance and 31% to 60% of the genetic variance in D9, D16, and D18 PF intake and compulsive-like eating, respectively (**Table 1**). Sex differences in bitter taste detection have been reported in humans [60] and in rats which can fluctuate across the estrous cycle [61] and could contribute to the phenotypic variance in females. Future genome-wide studies in female mice that seek to identify female-specific QTLs accounting for the unexplained variance in BE should also monitor and account for variance due to estrus cycle [62]. Indeed, our null result for females on chromosome 6 could potentially be explained by variance due to estrus cycle masking the genotypic effect on PF intake.

The *Tas2r* locus contains a cluster of 25 *Tas2r* genes that code for T2Rs that function as bitter taste receptors [33]. While there was no effect of Genotype on initial PF consumption at the *Tas2r* locus (**Fig.3D,E**), there was an effect at later time points where mice homozygous for D2J consumed more PF on D9 and D16 as well as greater compulsive-like PF consumption compared to mice homozygous for B6J (**Fig.3D-F).** Together, the results suggest that decreased sensitivity to bitter taste perception associated with the D2J allele could permit consumption of progressively larger amounts of sweetened PF. Or i.e., increased sensitivity to bitter taste associated with the B6J allele (reduced bitter consumption) could limit the escalation of PF consumption because of the bitterness itself and/or over-heightened the sweet perception of sucrose [63], thus limiting intake.

The association of the *Tas2r* locus with BE in our model has relevance for BE and eating disorders in humans [64]. Genetic influences on eating behaviors can potentially affect multiple biological processes that include taste processing and perception, palatability, liking/preference versus disliking/disgust/aversion, and/or potentially one or more neural circuits involved in energy homeostasis (hunger versus satiety) versus food motivation and reward [65, 66]. The genetic basis of eating disorders, and more specifically BE as a quantitative trait, is almost entirely unknown but polymorphisms in taste receptors could contribute to BE susceptibility. In support, a recent human genetic study identified a coding polymorphism within *TAS2R38* that decreased sensitivity to bitter taste was associated with increased disinhibition of eating behavior in women of Amish descent [67]. Additionally, in line with our current results, increased preference for sweetened food has been linked to decreased perception of bitter taste [68]. *TAS2R38* is a member of the bitter gene receptor family and accounts for up to 60-85% of the phenotypic variance in bitter sensitivity to synthetic pharmaceuticals [69].

Taste receptor polymorphisms are associated with the choice and amount of food that is consumed which can influence taste receptor physiology. Polymorphisms of several different *TAS2R* genes in humans are associated with an altered response to bitter taste stimuli and ingestive behaviors such as alcohol use and abuse [70, 71], vegetable consumption [72], and consumption of energy dense foods such as butter, beer, and cured meat [73]. Additionally, genetic variations in the sweet taste receptor *TAS1R2* are associated with dietary sugar intake in overweight and obese populations [71]. The type and amount of food consumed has been shown to be altered in BED compared to obese controls, in that subjects with BED choose to consume more energy-dense foods such as red meat and sweet foods such as ice cream [74]. Together, polymorphisms in taste receptors influence food selection which could increase risk for BE. In addition to taste receptor polymorphisms affecting eating behaviors, disordered eating behaviors such as cycles of BE and food restriction can alter taste receptor function, thus acting as a vicious cycle to facilitate disordered eating. Both AN and BN have been associated with reduced brain response to an aversive bitter stimulus [75] as well as reduced perception of bitter taste [76], and AN, BN, and BED are associated with reduced taste papillae [77]. Thus, the progressive increase in PF consumption in the bitter taste-resistant D2J strain at the *Tas2r* locus [26] could further alter taste receptor physiology and further add to the reduced genetic effect on bitter perception, thus, exacerbating BE.

In addition to the *Tas2r* locus, the D2J allele at the chromosome 11 locus (50 Mb) was associated with an enhancement of PF intake (D18) as well as the slope of escalation (**Fig.5G,H**). This locus accounted for 8% and 19% of the phenotypic variance as well as 8% and 26% of the genetic variance in D18 PF intake and slope of escalation, respectively. The effect was more pronounced in the males where 39% of the phenotypic variance and 73% of the genetic variance was captured by this locus (**Fig.6G-L**). Using mice comprising B6J and D2J alleles, this locus was previously associated with intravenous cocaine self-administration and differential expression of the gene *Cyfip2* [35], as well as methamphetamine-induced locomotor activity and the gene *Hnrnph1* [34, 36]. Thus, polymorphisms in either gene could potentially contribute to escalated consumption comprising BE, e.g., via modulation of the reward circuit function. In support, we previously mapped and validated *Cyfip2* as a major genetic factor underlying BE in a reduced complexity cross between B6 substrains [18].

There are several limitations to this study. Despite a careful genetic analysis and assessment of reasonable candidate loci, this study lacks the unbiased nature of a genome-wide QTL scan. While our F_2_ study was not well-powered to detect genome-wide significant
effects of QTLs of moderate effect size, it served as a reasonable preliminary investigation into the contribution of well-established taste and drug loci while at the same time assessing feasibility of conducting a genome-wide QTL analysis in a larger F_2_ cohort and employing the BXD genetic reference panel. Because of the small sample size, it is possible that our variance estimates are inaccurate. Hundreds of additional mice will need to be phenotyped (both PF and Chow pellets) and genotyped at a panel of markers to more accurately account for phenotypic variance and identify additional loci contributing to the various components of BE, in particular for females where a majority of the heritability remains unexplained.

To summarize, we identified a forward genetic model of BE that lays the groundwork and rationale for a future large-scale, well-powered forward genetic study, especially in female mice where more of the variance remains unexplained. The clear advantages offered by the genetic tools comprising alleles from the B6J and D2J parental strains, including the BXD RI panel [24], interval-specific congenics for fine mapping [78], and the wealth of phenotypic data on GeneNetwork [79, 80] ensure that a systems genetic approach with these powerful tools will likely yield new insight into the genetic, genomic, and neurobiological mechanisms of risk for BE as well as the neurobiological adaptations underlying the establishment of and recovery from BE which will inform new treatment strategies.

## ACKNOWLEDGEMENTS

These studies were supported by NIDA grants R21DA038738 (C.D.B.) and R01DA039168 (C.D.B.), We would like to acknowledge the Analytical Instrumentation Core at Boston University School of Medicine, including the Core Director, Dr. Lynn Lingyi Deng and the Lab Manager, Matthew Au.

## AUTHOR CONTRIBUTIONS

C.D.B. conceptualized and designed the experiments. R.K.B., J.C.K., J.L.S., and K.P.L. collected data from the food experiments. R.K.B. and J.C.K. analyzed the data. R.K.B. and C.D.B., and M.K.M. wrote the manuscript.

